# Quantifying ecosystem degradation via energetic deficits

**DOI:** 10.1101/2025.05.16.654496

**Authors:** Marco Tulio Angulo, Yitong Liu, Dan Morris, Serguei Saavedra

## Abstract

Ecosystem degradation is accelerating, threatening biodiversity and human economies. Quantifying this degradation relative to a reference baseline remains a key challenge. Yet, this quantification is essential to provide the necessary resources to regenerate ecosystems. We address this challenge by studying the energetics of population dynamics, where energy inputs such as sunlight and rainfall are transformed into other forms of energy embodied as biomass. This approach provides a unifying framework to quantify degradation across diverse ecosystems via the *energetic ecodeficit*: the extra energy supply the degraded ecosystem would require to regenerate to a reference state. This energetic deficit can then be translated into a *monetary ecodeficit*, representing the economic cost of regeneration, considering the local cost of embodied energy. Our analysis reveals nonlinear relationships between biomass loss and energetic ecodeficit, including points of no return beyond which restoration becomes economically infeasible. We showcase our methodology using the current mammal deficits across the African savanna, finding that the monetary ecodefict is about 100 times larger than current estimates. Our work demonstrates how integrating energetic principles into conservation planning can transform ecosystem valuation globally, linking ecosystems and economies sustainably.

## 1 Introduction

Ecosystems are being degraded at an alarming rate, largely due to human activities like massive habitat destruction and rapid climate change (Dasgupta and Levin, 2023, Dirzo et al., 2014, Scheffers et al., 2016). Quantifying ecosystem degradation is therefore essential for achieving planetary sustainability (Strassburg et al., 2020) and for measuring the loss of natural capital in terms of the biological elements that support life’s adaptation (Ouyang et al., 2017). However, quantifying the degradation of an ecosystem relative to a reference state —its “ecodeficit”— has proven difficult (Moreno-Mateos et al., 2017). One major challenge is that ecosystems are extremely diverse in terms of components (e.g., species) and environmental conditions (e.g., habitat, geography, climate, etc.). This diversity makes it difficult to identify a “common currency” to quantify degradation across ecosystems as different as the Amazon rainforest and coral reefs. It is also often difficult to determine what the reference state of a degraded ecosystem should be, particularly when the exact mechanisms of degradation remain unclear. (Moreno-Mateos et al., 2017, Toit et al., 2003). Another important challenge is that it remains unclear how to quantify ecosystem degradation in a way that can assist their regeneration (Odum, 1969).

To address these challenges we draw upon the fundamentals of ecosystem energetics (Lindeman, 1942, Odum and Barrett, 2005, Pielou, 2001), proposing energy as the common currency needed to quantify ecodeficits across diverse ecosystems. From an energetic perspective, all ecosystems can be conceptualized as processes that transform some energy supply, such as solar radiation, into biomass via photosynthesis and subsequent trophic interactions (Drake et al., 2007). The transformation from energy to biomass is mediated by the energy use of organisms and their interactions (Barneche and Allen, 2018, Campillay-Llanos et al., 2022, Thompson and Townsend, 2005). We postulate that disturbances such as deforestation, pollution and habitat degradation perturb ecosystems by distorting their energetic efficiency (Tabi et al., 2023, Warren and Spencer, 1996). In degraded ecosystems, the energy-to-biomass transformation is less efficient, resulting in lower biomass levels than reference values. This observation allows us to define the *energetic ecodeficit* as the additional energy supply that a degraded ecosystem would require to regenerate to a reference state. The reference state can be defined as a historical or theoretical baseline with intact energy flows. Energy also serves as an instrumental bridge to economics (Odum, 1996), allowing us to translate the energetic ecodeficit into a *monetary ecodeficit* that considers the economic cost of supplying this additional energy. This monetary valuation can inform policy decisions, prioritize restoration efforts, and integrate ecological and economic considerations into sustainability planning.

The rest of this paper is organized as follows. We first formally define energetic ecodeficits. Next, we use model ecosystems to analyze the energetic ecodeficit associated with different perturbations. Model ecosystems enable the controlled perturbations necessary to study ecodeficits in detail (manipulations that would be impossible in real ecosystems). We find non-linear ecodeficit responses to perturbations, with singularities beyond which regeneration becomes infeasible. We then study the ecodeficit in real-world ecosystems using the African savanna as a case study. Ecosystems in the African savanna have experienced various forms of degradation, from soil erosion to population declines, making it difficult to quantify how much they have actually degraded (Toit et al., 2003). We demonstrate how the ecodeficit can be estimated by combining empirical data (i.e., biodiversity census), estimates for intact energy flows, and theoretical expectations for energy use of individuals. We end by discussing the implications of our findings for biodiversity conservation and ecosystem management.

## 2 Materials and Methods

To quantify energetic ecodeficits, consider an ecosystem with *m* different sources of energy supplied at rates 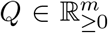 (units of energy/area/time). These could be solar radiation, chemical energy from rainfall, or other relevant sources. Let the vector 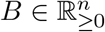 denote the ecosystem’s biomass densities (units of mass/area), where *n* ≥ 1 represents the number of distinct components (e.g., species, trophic levels, functional groups, etc.). The transformation from energy supply *Q* to biomass *B* is mediated by the ecosystem’s population dynamics, characterized by a vector *θ* ∈ ℝ^*p*^ of ecological parameters (e.g., uptake rates, mortality rates, migration rates, metabolic rates, among others).

Let *θ*^ref^ ∈ ℝ^*p*^ and 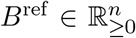 denote the reference values that the ecological parameters and biomass densities take before perturbations, respectively. These reference values are inter-dependent, as biomass densities depend on both the ecological parameters and the supplied energy. Such interdependence can be formally expressed as *f (B*^ref^, *θ*^ref^, *Q)* = 0, where *f* : ℝ^*n*^ *×* ℝ^*p*^ *×* ℝ^*m*^ → ℝ^*n*^ represents the functional relationship between variables. For example, in the classic Lotka-Volterra equations for ecological populations, the Volterra principle guarantees that the functional form *f* of the population dynamics satisfies this condition with *B*^ref^ the (time) average biomass densities (Hofbauer and Sigmund, 1998, Theorem 5.2.3). In our results, we show that the specific functional form of *f* can be deduced from principles of energy conservation.

Human activities often perturb ecosystem by altering their ecological parameters, such as affecting species interactions or metabolic rates (Bauer et al., 2015, Nicholson et al., 2023, Tabi et al., 2023, Warren and Spencer, 1996). We represent this perturbation via a multi-variable map *β* : ℝ^*p*^ → ℝ^*p*^ that changes the ecological parameters from a reference value *θ*^ref^ to the new perturbed value *θ*^pert^ = *β*(*θ*^ref^) ∈ ℝ^*p*^. For simplicity, we focus on perturbations that scale each ecological parameter by a factor *β*_*i*_ ≥ 0 (i.e., 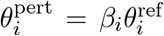). When *β*_*i*_ *>* 1 the perturbation increases the *i*-th ecological parameter, when *β*_*i*_ = 1 the perturbation leaves it unchanged, and when *β*_*i*_ *<* 1 the perturbation decreases it. The perturbation leads to a change in the biomass densities to a new perturbed value *B*^pert^ ∈ ℝ^*n*^. For example, in Fig. 1a the perturbation reduces all densities compared to the reference values.

**Figure 1:**
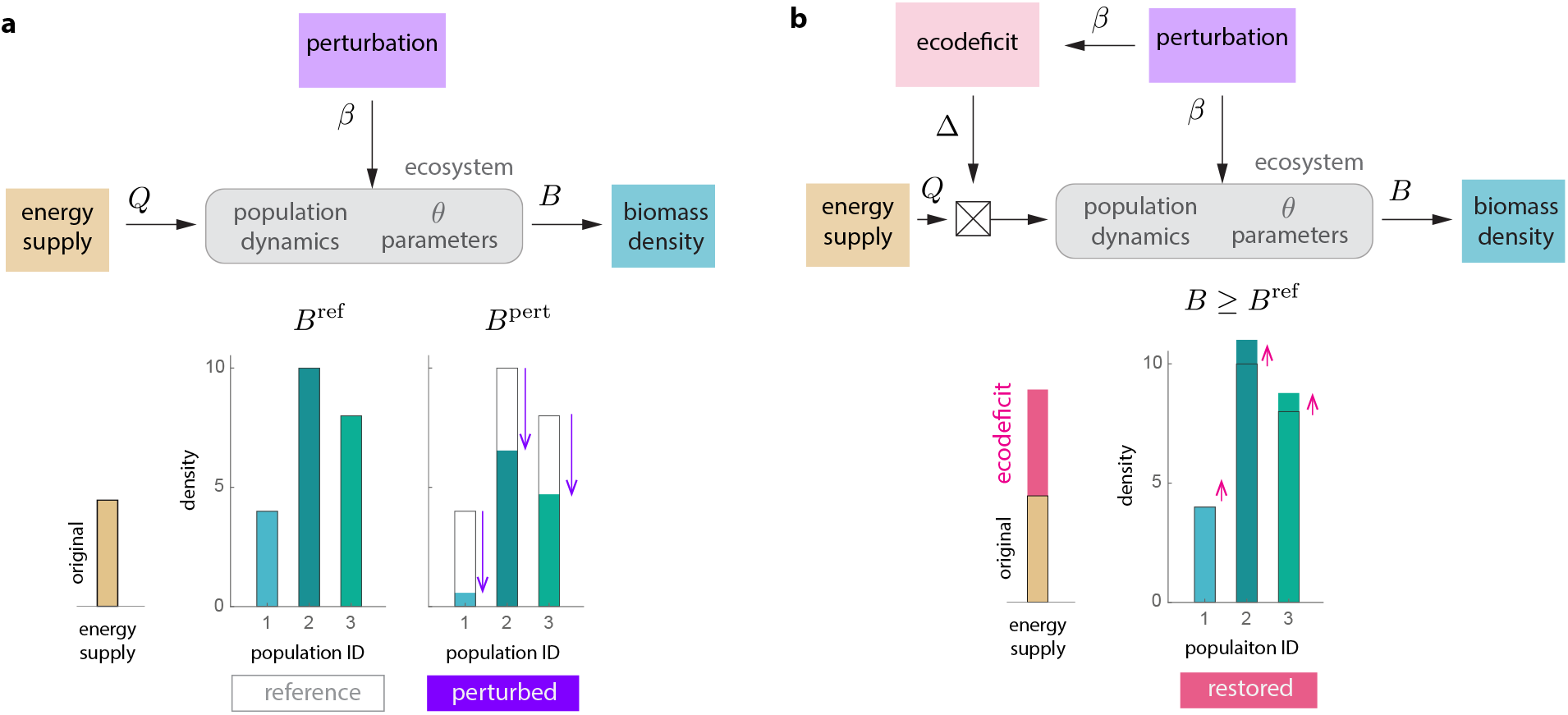
Quantifying the energetic ecodeficit of degraded ecosystems. Panels illustrate the concept using an hypothetical ecosystem of *n* = 3 populations. **a**. Let the vector 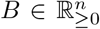 represent the biomass density of each population. The observed biomass densities depend on the energy input *Q* ∈ ℝ^*m*^ supplied to the ecosystem, and the population dynamics characterized by a vector *θ* ∈ ℝ^*p*^ of ecological parameters (e.g., uptake rates, mortality rates, metabolic rates, etc.). Let *θ*^ref^ denote the value of the ecological parameters without or before the perturbation, resulting in a vector 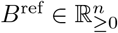 of reference biomass densities. Human activities perturb the ecosystem, changing its ecological parameters to *θ*^pert^ = *β*(*θ*^ref^) ∈ ℝ^*p*^ where the map *β* : ℝ^*p*^ → ℝ^*p*^ characterizes the perturbation. In response, the biomass densities are perturbed to *B*^pert^. In the example shown in the panel, the perturbation reduces the density of all populations. **b**. Let ∆ : ℝ^*m*^ → ℝ^*m*^ be a diagonal map representing a change to the ecosystem energy supply. Given the perturbed ecosystem, the energetic ecodeficit factor is defined as the minimum change ∆(*Q*) to the ecosystem supply *Q* such that the resulting population densities *B* satisfy *B ≥ B*^ref^ entry-wise. That is, the minimum change to the ecosystem energy supply to regenerate its biomass density to at least its reference values.

To calculate the energetic ecodeficit, consider rescaling each energy supply *Q*_*j*_, *j* = 1, *· · ·*, *m*, by a constant adimensional factor ∆_*j*_ ≥ 0. We use the notation 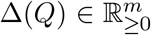 to denote vector of all rescaled energy supplies. The energetic ecodeficit is then defined as the the solution to the following optimization problem:

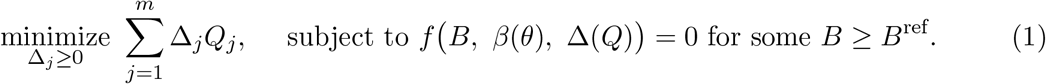

The ecodeficit represents the minimum change ∆ such that, when applied to the energy supply *Q* of the ecosystem, results in biomass densities that meet or exceed the reference densities (Fig. 1b). We are seeking the minimum additional energy required to restore the ecosystem. Mathematically, this can be interpreted as a conjugacy-like condition *f(B, β*(*θ*), ∆(*Q*)) = 0, determining the energetic ecodeficit ∆(*Q*) associated to the perturbation *β*(*θ*). When no amount of additional supply can restore the ecosystem, the optimization problem of Eq. (1) has no solution, and we define the ecodeficit as ∆_*j*_ = ∞. Conversely, an ecodeficit *<* 1 means the perturbation improved the ecosystem’s energy efficiency, allowing it to reach the reference densities using a lower energy supply.

From the solution to Eq. (1) we can derive three ecodeficit metrics. First, the *ecodeficit factor* 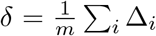 quantifying, on average, how many times each energy supply must be multiplied to regenerate the ecosystem to reference values. Note that *δ* = ∆ for *m* = 1 energy supply. Second, the *energetic ecodeficit* ∆_*E*_ = ∑ _*i*_(∆_*i*_ − 1)*Q*_*i*_ quantifies the additional energy supply that the ecosystem requires for regeneration (units energy/area/time). Third, the *monetary ecodeficit* ∆_*M*_ = *k*∆_*E*_ represents an economic quantification of the cost of supplying the energetic ecodeficit (units money/area/time), with *k* ≥ 0 an energy-to-money conversion factor (Fig. 1d). We assume this factor is country-specific, and it can be approximated as the total embodied energy used by the country in a given year divided by its Gross Domestic Product (GDP) in that year (Odum, 1996, Sweeney et al., 2007). Note that different from absolute energy, embodied energy (also known as *emergy*) quantifies the cumulative energy that was needed to transform into a given current state of energy (Odum, 1996). Embodied energy is measured in terms of solar energy units, and it is equivalent to the input energy *Q* obtained by ecosystems needed to transform into different biomass forms (Odum and Barrett, 2005).

## 3 Results

### Energetic ecodeficits in model ecosystems

We focus our study on trophic chains with *n* = 3 levels (Hofbauer and Sigmund, 1998, Logofet, 1993, Loreau, 2010). Each trophic level specifies a discrete supply level from one form of energy to a different one, rather than categorizing individual populations (MacArthur, 1955, Odum and Barrett, 2005). Specifically, we consider primary producers such as plants (level 1), primary consumers such as herbivores (level 2), and secondary consumers such as carnivores (level 3). Each level *i* has a characteristic *energy content µ*_*i*_ *>* 0 per unit biomass (units energy/mass) and receives energy from the previous trophic level. An implicit basal resource (level 0) receives a constant external energy supply *Q* (units of energy/area/time).

We derived the energy dynamics of the trophic chain from an energy conservation argument, resulting in a system of *n* = 3 differential equations (Methods and Supplementary Note 1.1). These equations share the same structure as classic models derived from population-based arguments (Hofbauer and Sigmund, 1998, Logofet, 1993, Loreau, 2010). However, they are parametrized differently, using three sets of parameters describing the flow of energy within each trophic level (Fig. 2a). First, the *assimilation efficiency f*_*i*_ ∈ [0, 1) (adimensional) is the fraction of the energy available to trophic level *i* that is assimilated. Second, the steady-state *production efficiency J*_*i*_ ∈ [0, 1) (adimensional) is the fraction of the assimilated energy used to generate new biomass. Finally, the mass-specific *respiration rate d*_*i*_ ≥ 0 (units of energy/mass/time) is the rate per unit biomass at which energy is used for all activities other than biomass production (e.g., maintenance, competition, etc.) Typical values for these parameters are shown in Supplementary Table S1.

**Figure 2:**
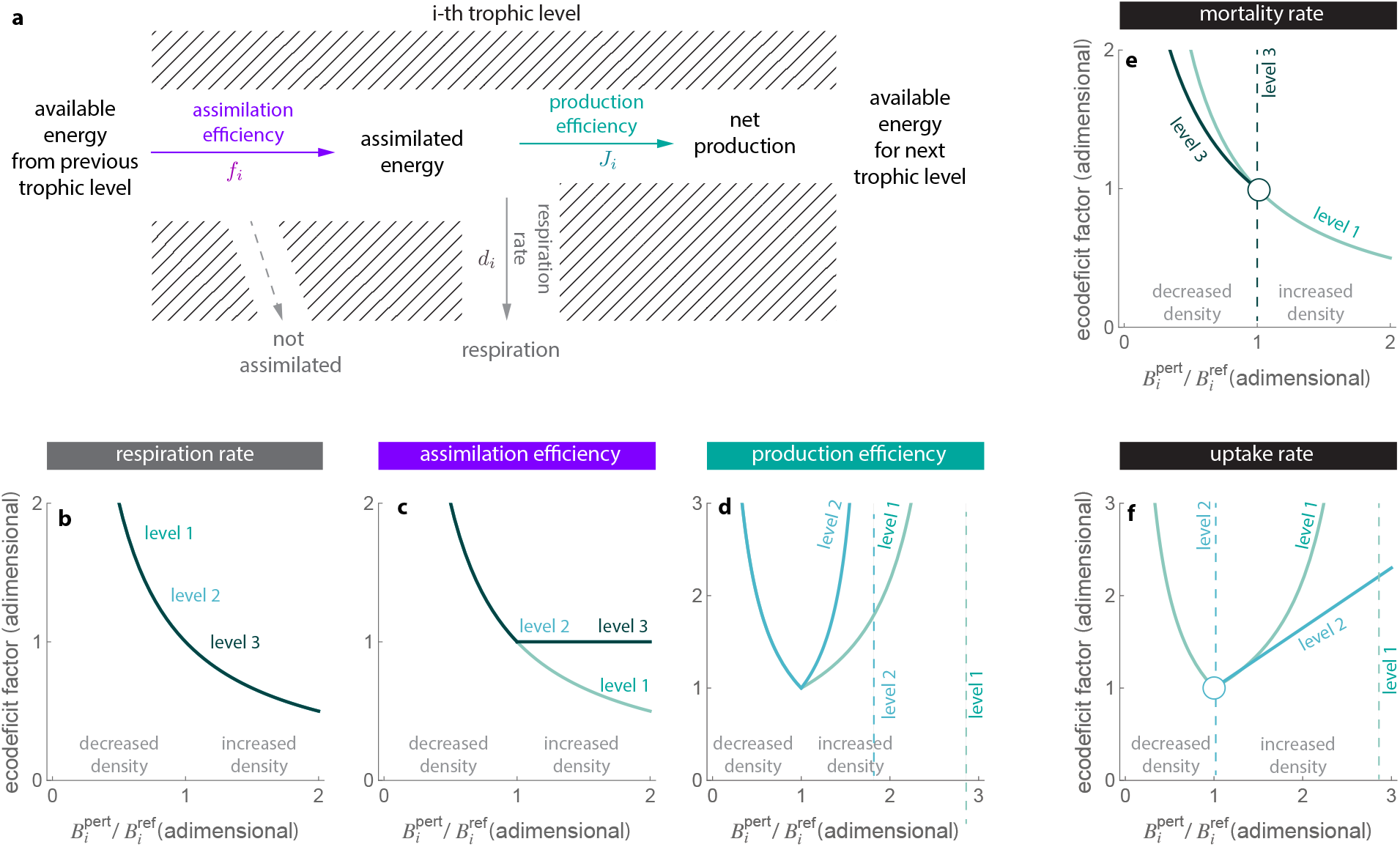
Energetic ecodeficit in model ecosystems. Results are for a mathematical model of a trophic chain with *n* = 3 levels derived from energy conservation arguments (Methods). Level 1 corresponds to primary producers such as plants, level 2 to secondary producers such as herbivores, and level 3 to tertiary producers such as predators. **a**. The model is characterized by three energy-based parameters that describe energy flow within each trophic level *i* = 1, 2, 3. The available energy supply at the input of each trophic level represents its energy intake. For strict autotrophs, this energy source is light, while for strict heterotrophs, it is organic food. The assimilation efficiency *f*_*i*_ ∈ [0, 1) (adimensional) is the fraction of the energy available in the previous trophic level that is assimilated. Energy that is not assimilated may include food egested without being metabolized, or light that passes through vegetation without being fixed. The production efficiency *J*_*i*_ ∈ [0, 1) (adimensional) is the fraction of the assimilated energy used for producing new or different organic matter. This represents primary production in plants and secondary production in animals, and it constitutes the energy available for the next trophic level. Respiration accounts for the energy used for all activities other than production. The biomass-specific respiration rate *d*_*i*_ *≥* 0 (energy/mass/time) characterizes the rate at which energy is used for respiration per unit biomass. **b-d**. The ecodeficit factor ∆_*i*_(*β*) associated with perturbing an energetic parameter *θ*_*i*_ at trophic level *i*, where the parameter changes from *θ*_*i*_ to *βθ*_*i*_ for some *β >* 0 characterizing the perturbation magnitude (see Table 1 for expressions). Panels illustrate the ecodeficit factor ∆_*i*_ when the perturbation magnitude *β* is written as a function of the ratio 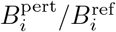 of perturbed to reference biomass densities. Labels denote different trophic levels. Dashed lines indicate asymptotes where the ecodeficit factor approaches infinity. **e-f**.The ecodeficit ∆_*i*_ associated with perturbing population-based parameters. Panel e shows the results when perturbing each trophic level’s mortality rate *m*_*i*_ *>* 0 (number of deaths per unit time). Panel f shows the results when perturbing each trophic level’s uptake rate *α*_*i*_ (the rate at which the population at trophic level *i* utilizes resources from the previous trophic level per unit biomass of level *i*). Circles represent discontinuities where the ecodeficit abruptly changes to infinity. Dashed lines represent asymptotes where the ecodeficit grows to infinity.

Our energy-based model allows us to derive analytical expressions for the ecodeficit by solving Eq. (1) (Methods and Supplementary Note 1.3). Table 1 summarizes the resulting expressions. These expressions can be more broadly interpreted by visualizing the ecodeficit factor ∆_*i*_ as a function of the ratio 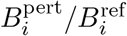 of perturbed to reference biomass densities. This analysis reveals three distinct types of ecodeficit responses (Fig. 2b-d).

**Table 1:** Ecodeficit associated with perturbing trophic chains. The first column indicates the perturbed parameter. We represent the perturbation to the parameter *θ* as *θ ↦ βθ* where *β >* 0 characterizes the perturbation magnitude (see Main Text). The second column shows that we can map the magnitude *β* of the perturbation to the ratio 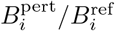 of perturbed to reference (i.e., unperturbed) biomass densities. Cells show how *β* is written as a function of this ratio for perturbations to different parameters. The last three columns show the ecodeficit factor ∆_*i*_ = ∆_*i*_(*β*) associated with perturbing each parameter in each trophic level.

| perturbed parameter | perturbation $\beta$ as biomass change | ecodeficit level 1 ($\Delta_1$) | ecodeficit level 2 ($\Delta_2$) | ecodeficit level 3 ($\Delta_3$) |
| --- | --- | --- | --- | --- |
| respiration rate ($d_i$) | $\frac{1}{B_i^{\text{pert}}/B_i^{\text{ref}}}$ | $\max\{1, \beta\}$ | $\max\{1, \beta\}$ | $\max\{1, \beta\}$ |
| assimilation efficiency ($f_i$) | $B_i^{\text{pert}}/B_i^{\text{ref}}$ | $\frac{1}{\beta}$ | $\max\left\{1, \frac{1}{\beta}\right\}$ | $\max\left\{1, \frac{1}{\beta}\right\}$ |
| production efficiency ($J_i$) | $\frac{1 - (1 - J_i)B_i^{\text{pert}}/B_i^{\text{ref}}}{J_i}$ | $\max\left\{\frac{1}{\beta}, \frac{1 - J_1}{1 - \beta J_1}\right\}$ | $\max\left\{\frac{1}{\beta}, \frac{1 - J_2}{1 - \beta J_2}\right\}$ | — |

First, perturbing the respiration rates *d*_*i*_ results in an ecodeficit factor that is inversely proportional to the ratio 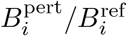 and that is identical across all trophic levels (Fig. 2b). When respiration increases, causing a decline in biomass density, the ecodeficit ∆_*i*_ also increases. In particular, ∆_*i*_ → ∞ as 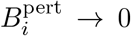. Conversely, when respiration decreases, resulting in an increase in biomass density, the ecodeficit factor falls below 1. This implies that the ecosystem improves its energetic efficiency as 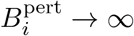.

Second, perturbing the assimilation efficiencies *f*_*i*_ results in different ecodeficit responses for different trophic levels (Fig. 2c). For primary producers (level 1), the ecodeficit factor ∆_1_ mirrors the response observed for respiration rates. However, for higher trophic levels (levels 2 and 3), the ecodeficit factors are always greater than one (i.e., ∆_*i*_ ≥ 1). This implies that while increasing the assimilation efficiency of primary producers can reduce the ecodeficit below 1, doing so for higher trophic levels does not have the same effect. This difference arises because primary producers directly utilize the external energy supply, while higher trophic levels rely on energy transfer from lower levels. Therefore, increasing the assimilation efficiency of higher trophic levels can enhance their energy uptake but does not necessarily improve the overall energy efficiency of the ecosystem. Note also that perturbations to the respiration rates or assimilation efficiencies that increase biomass density do not degrade the ecosystem because their ecodeficits satisfy ∆_*i*_ ≤ 1.

Finally, perturbing production efficiencies *J*_*i*_ yields the most complex ecodeficit response, where both increases and decreases in the biomass densities lead to a higher ecodeficit factor (Fig. 2d). This counterintuitive behavior occurs because changes in production efficiency at one trophic level can indirectly affect energy flow and respiration at other trophic levels. For instance, increasing the production efficiency of herbivores may lead to greater energy consumption and respiration by carnivores, potentially increasing the overall ecodeficit. Simply increasing the energetic efficiency somewhere in the ecosystem will not always decrease the ecodeficit. Furthermore, the ecodeficit factor has an asymptote at 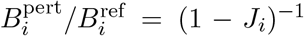 at which the ecodeficit becomes infinity. This asymptote, which appears first for secondary consumers (level 2) and then for primary producers (level 1), highlights the potential for non-linear and even catastrophic ecodeficit responses to seemingly minor changes in energy flow parameters.

### Population-level parameters produce complex energetic ecodeficits

Classical trophic chain models, not derived from energy-based arguments, use population-based parameters such as *mortality rates m*_*i*_ (units of 1/time) and *uptake rates α*_*i*_ (units of 1/time *×* 1/units of *B*_*i*_). To link our energy-based framework with these classical models, we mapped these population-based parameters as functions of our energy-based parameters (Supplementary Note 1.2):

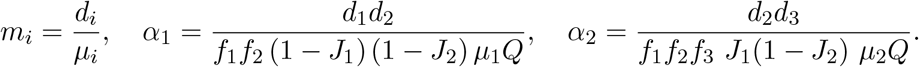

Using the above equivalence, we can investigate how perturbations in population-level parameters affect the energetic ecodeficit(Supplementary Note 1.3).

Our analysis reveals that perturbing population-level parameters leads to more complex ecodeficit responses compared to perturbing energy-based parameters (Fig. 2e-f). Specifically, the ecodeficit factor exhibits asymptotes where the ecodeficit factor grows to infinity (dashed lines in Fig. 2e-f) and discontinuities where the ecodeficit factor abruptly jumps to infinity (circles in Fig. 2e-f). For instance, increasing the mortality rate *m*_3_ of the last trophic level results in an infinite ecodeficit (Fig. 2e). Similarly, decreasing the uptake rate *α*_2_ equally results in an infinite ecodeficit (Fig. 2f). These extreme responses occur because such perturbations can drive the biomass density of the middle trophic level below their reference state, and no amount of additional energy can restore the system.

### Energetic ecodeficits in the African savanna

To demonstrate the applicability of our framework to real-world ecosystems, we study the energetic ecodeficits associated with the decline of large mammal populations in the African savanna (Ripple et al., 2015). This decline is a strong indicator of degradation in these ecosystems (Craigie et al., 2010), as large herbivores and carnivores play key roles in various ecological processes such as structuring vegetation, nutrient cycling, seed dispersal, and top-down control of prey populations (Estes et al., 2011, McNaughton, 1985, Sinclair et al., 2003). The African savanna can be conceptualized as a trophic chain, with solar radiation and rainfall serving as the primary energy sources (McNaughton, 1985).

Our analysis focuses on 18 ecological reserves across the African savanna, for which detailed biodiversity census data are available (Hatton et al., 2015) (see Table 2 for a summary of the data). These reserves exhibit diverse geographic, climatic, and ecological characteristics (Fig. 3a). For instance, average annual precipitation varies significantly across reserves, ranging from 207 mm in the Kalahari to 1894 mm in the Masai Mara. The composition of large mammal communities also differs substantially, with the number of herbivore species ranging from 5 in Katavi to 23 in Kruger National Park, and the number of carnivore species ranging from 2 in Gonarezhou and Queen Elizabeth National Park to 5 in the Kalahari (Fig. 3b). This diversity highlights the ecological variability of savanna ecosystems even within a broadly similar biome.

**Table 2:** Summary of the African data used for the analysis. We studied 18 reserves compiled from previous work (Hatton et al., 2015). Data for Net Primary Productivity was obtained from FAO’s WaPOR database (Fao, 2025). We considered a square of 200 *km*^2^ centered in that location. Then we used FAO’s WaPOR database to retrieve the average NPP (layer “L1-NPP-D”) over that square with data from the first to the last census. The average NPP and average *B*_2_ (prey) *B*_3_ (predator) of the reservation is taken to be the temporal average of this data.

| **Reserve**<br>**(name)** | **census**<br>**(year)** | **average area**<br>**(km<sup>2</sup>)** | **average NPP**<br>**(kcal/m<sup>2</sup>/day)** | **average B2**<br>**(kg/km<sup>2</sup>)** | **average B3**<br>**(kg/km<sup>2</sup>)** |
| --- | --- | --- | --- | --- | --- |
| Amboseli | 2007 | 390 | 8.96 | 7103.13 | 63.18 |
| Etosha | 1926,1998 | 16280 | 0.93 | 198.76 | 4.65 |
| Gonarezhou | 2009 | 5053 | 7.83 | 479.70 | 10.16 |
| Hluhluwe | 1982,2000 | 960 | 9.88 | 4397.99 | 37.53 |
| Hwange | 1973,1996 | 14651 | 9.78 | 512.35 | 8.59 |
| Kalahari | 1979 | 9590 | 0.80 | 257.88 | 3.82 |
| Katavi | 2009 | 4471 | 12.31 | 2795.73 | 14.76 |
| Kidepo | 2009 | 1442 | 9.86 | 1423.62 | 14.43 |
| Kruger | 1964,1975,1984,1997,2009 | 18989 | 7.42 | 1361.44 | 20.38 |
| Lake Manyara | 1970,1992 | 110 | 9.99 | 8076.97 | 56.57 |
| Masai | 1992,2003 | 1500 | 15.38 | 4391.58 | 67.10 |
| Mkomazi | 1970,1970 | 3245 | 8.33 | 283.13 | 5.03 |
| Nairobi | 1966,1976,2002 | 117 | 5.08 | 3149.05 | 39.38 |
| Ngorongoro | 1965,1978,1988,1997,2004 | 260 | 14.73 | 13352.81 | 91.52 |
| Queen | 2009 | 1978 | 12.49 | 2414.02 | 18.07 |
| Selous | 1994 | 43626 | 11.66 | 2079.28 | 31.64 |
| Serengeti | 1971,1977,1986,1993,2003 | 25000 | 12.35 | 2158.13 | 26.85 |
| Tarangire | 1962,1962,2000 | 2850 | 10.90 | 1684.62 | 16.94 |

**Figure 3:**
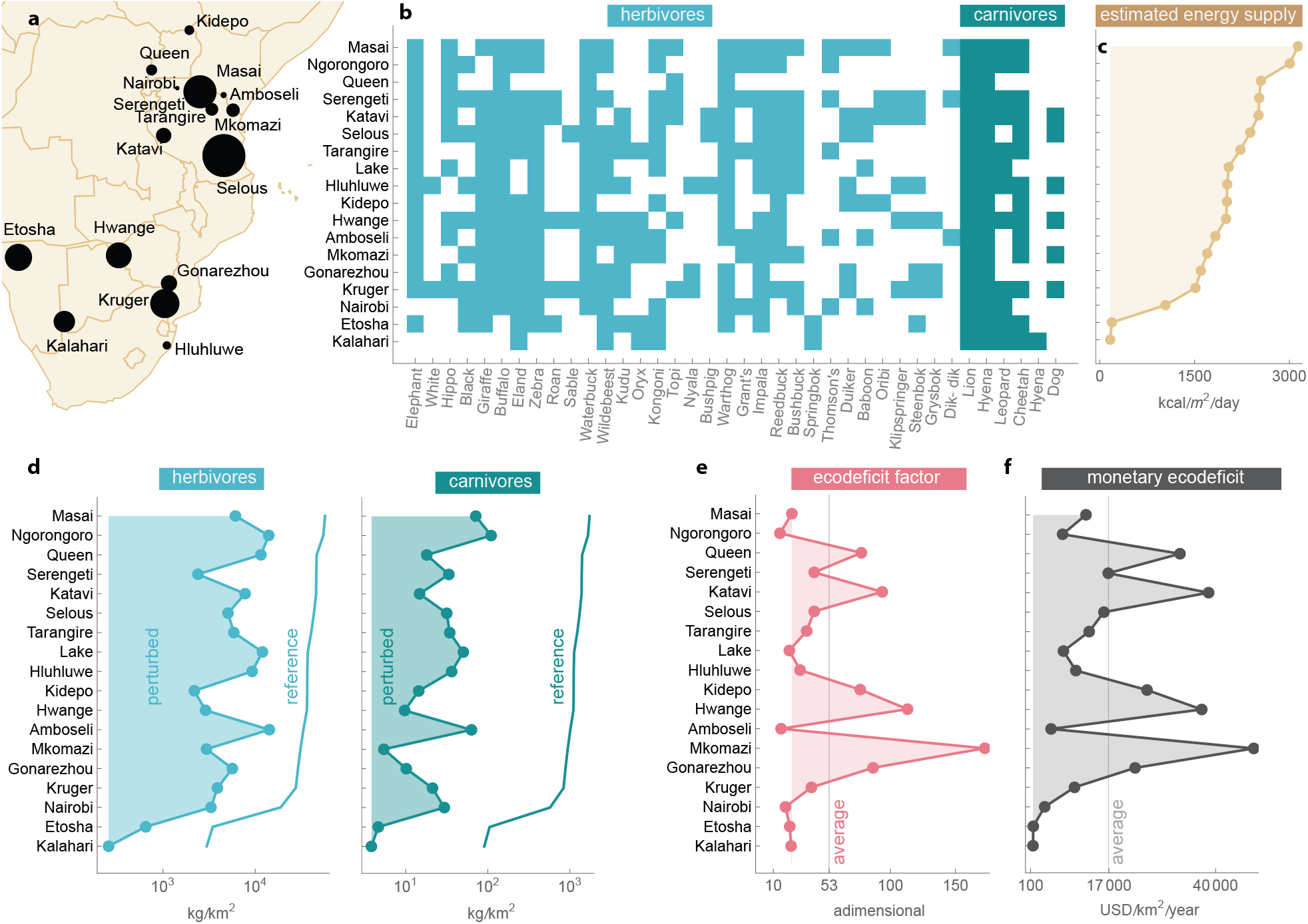
Ecodeficit associated with the decline of large mammals in the African savanna. **a**. We focus our study on 18 broadly distributed reserves where census data is available. Circle size is proportional to the reserve’s area, ranging from 110 (Lake) to 46,626 km^2^ (Selous). Table 2 provides a summary of the data. **b**. Reserves span various geographic and climatic conditions, for example, with different precipitation (from 207 to 1894 mm per year) and biome (from semi-arid sandy savannas to vast grasslands). These differences produce different ecosystems with diverse compositions of herbivores and carnivores. The panel illustrates this point, showing which of the 32 herbivores and 6 carnivores are present in each reserve. Reserves are ordered from high (Masai) to low (Kalahari) estimated energy supply. **c**. We used satellite data of Net Primary Productivity (NPP) at the location of each reserve to estimate its energy supply 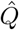 (see Main Text for details). Ecosystem differences are well explained by energy supply differences (e.g., reserves with higher energy supply tend to have higher diversity). **d**. We used previously compiled census data for each reserve (Hatton et al., 2015) to estimate the perturbed biomass density 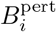 of herbivores (*i* = 2) and carnivores (*i* = 3) in each reserve (filled areas). We estimated the reference biomass densities 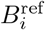 assuming the unperturbed ecosystem has the same energy supply but intact energy flows and respiration mainly driven by the metabolic needs of organisms (lines). **e**. The ecodeficit factor ∆ in each reserve is the number of times we need to multiply the energy supply to regenerate the ecosystem above its reference biomass densities. Eq. (2) of the main text shows how to estimate the ecodeficit factor from the above data and assumptions. The estimated ecodeficit factors are heterogeneous, indicating significant differences in ecosystem degradation across reserves. The Ngorongoro Crater reserve in Tanzania has the smallest ecodeficit factor (∆ = 15.1), which interestingly started being protected by restricting hunting very early in 1921. The Mkomazi Game reserve in Tanzania has the larger ecodeficit factor (∆ = 172.6), which historically has faced significant challenges with cattle. We find an average ecodeficit factor ∆ = 53 across reserves. **f**. We calculated the monetary ecodeficit using the most recent average conversion *k* from embodied energy (emergy) to USD (Eme, 2025). On average across African reserves, we find an average monetary ecodefict of about 17, 000 USD/km^2^/year, with the highest ecodeficit of 48, 234 USD/km^2^/year occurring for Mkomazi (9, 590 km^2^) and the lowest of 615 USD/km^2^/year occurring for Kalahari. Notice that Kalahari and Etosha are regions with low energy input and low ecodeficit, revealing that even small energy supplies can help to restore these ecosystems.

To estimate the ecodeficit for each reserve, we make the following assumptions about the unperturbed state of the ecosystem: (i) the energy flows are intact, corresponding to the upper ranges of assimilation 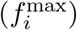 and production 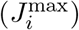 efficiencies; (ii) the external energy supply is the same in the perturbed and reference states; and (iii) respiration is primarily driven by the metabolic needs of the organisms. Under these assumptions, the ecodeficit factor ∆ in a reserve is (Methods):

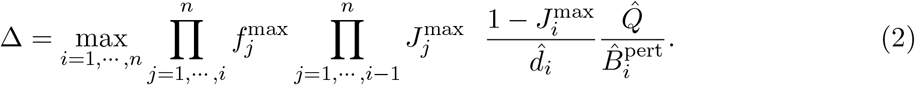

We use field-work data from various ecosystems to estimate 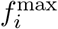 and 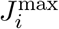 (see Supplementary Table S1 and (Odum and Barrett, 2005, pp. 110)). Based on these estimates, we can use assumption (ii) to estimate the energy supply as 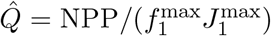 using satellite data on net primary productivity (NPP) from FAO’s WaPOR database (Fao, 2025).

The estimated energy supply 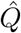 varies substantially across reserves, spanning a 19-fold range (Fig. 3c). The lowest energy supply is observed in the Kalahari, a semi-arid sandy savanna, with an estimated 164 kcal/m^2^/day (about 8 Watts per m^2^). In contrast, the highest energy supply is found in the Masai Mara, a vast grassland savanna, with an estimated 2225 kcal/m^2^/day (about 107 Watts per m^2^). Powering a single 100W light bulb requires about 12 square meters in the Kalahari but only one square meter in the Masai. This considerable variation in energy supply across reserves highlights energy as an adequate summary of the many environmental factors influencing these ecosystems.

Following previous work on biomass estimation in these reserves (Hatton et al., 2015), we estimate the perturbed biomass density from census data as 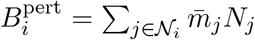. Here, *𝒩*_*i*_ is the set of all populations belonging to trophic level *i*, 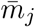 is the average body size of individuals of population *j* (units of kg), and *N*_*j*_ is the number of individuals of population *j* per area reported in the biodiversity census (units of 1/km^2^). Overall, reserves with higher energy supply tend to exhibit higher biomass densities of herbivores and carnivores (Fig. 3d).

From assumption (iii), we can estimate the mass-specific respiration rate *d*_*i*_ using allometric scaling relationships based on Kleiber’s and Damuth’s laws (Damuth, 1987, Kleiber, 1932). Kleiber’s law relates the field metabolic rate to body mass, while Damuth’s law connects population density to body mass. Combining these laws yields the estimate 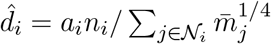 (Methods and Supplementary Note S2.1). Here, *n*_*i*_ is the number of populations in trophic level *i*, and *a*_*i*_ is the expected field metabolic rate of an individual of trophic level *i* with unit body size (Supplementary Table 2).

Using the estimated energy supply, biomass densities and respiration rates, we calculate the ecodeficit factor ∆ for each reserve using Eq. (2). Figure 3e shows the ecodeficit factor listed in ascending order of input energy. Ecodeficit factors vary considerably across reserves, ranging from ∆ = 15.1 in Ngorongoro Crater to ∆ = 172.6 in Mkomazi Game Reserve. This variation indicates a wide range of degradation across the savanna ecosystems. Interestingly, the ecodeficit does not exhibit a simple monotonic relationship with energy supply, suggesting that factors beyond energy availability contribute to ecosystem degradation.

To translate the energetic ecodeficit into a monetary value, we use the conversion factor *k* that relates embodied energy (solar equivalent Joules) to USD (Fig. 3f). We use the most recent estimate of this conversion factor, *k* = 9.24 *×* 10^12^, which represents an average value across Namibia, Kenya, Zimbawe, South Africa, Uganda, and Tanzania (Eme, 2025). Normalizing across reserves as a function of their area, we found an average monetary ecodeficit of 16, 401 USD/km^2^/year for Africa. Extrapolating this value to the entire African savanna (13.5 *×* 10^6^ km^2^) yields about 0.22 trillion USD per year. The largest monetary ecodeficit across reserves occurs in Mkomazi Game Reserve, with an estimated 48, 234 USD/km^2^/year, equivalent to an annual loss of 0.2 billion USD for this reserve alone. These ecodeficits quantify the natural capital that is lost every year per region or reserve.

## 4 Discussion

The accelerating degradation of ecosystems poses a significant threat to biodiversity and human well-being, with potentially irreversible consequences on people, species, and economies (Moreno-Mateos et al., 2017, Scheffers et al., 2016). Yet, given the inherent complexity of ecosystems (Moreno-Mateos et al., 2017), it remains unclear and debated if and how shall we quantify ecosystem deficits in a way that could de-accelerate anthropogenic effects. While prominent work has used econometric approaches and meta-data analyses to address this gap (Alkemade et al., 2009, England, 2023, Museum, 2023, Stanford University and the Royal Swedish Academy of Sciences., 2024), here we introduced a general theoretical framework rooted in the fundamental principles of energy. Energy is a common currency of life, driving virtually all biological processes (Lotka, 1922, Odum, 1996). Following this premise, the notion of energetic ecodeficit we introduce quantifies the additional energy that a perturbed ecosystem requires to regenerate to a reference state. This notion is closely related to concepts of available storage and required supply introduced in dynamical systems theory (Willems, 1972). The energetic ecodeficit can then be translated into a monetary ecodeficit, providing an economic measure of the restoration costs. Our framework aligns with the growing recognition of how perturbations can fundamentally alter the energetics of ecosystems (Barneche and Allen, 2018, Campillay-Llanos et al., 2022, Tabi et al., 2023, Thompson and Townsend, 2005, Warren and Spencer, 1996).

Using a trophic chain model ecosystem, we demonstrated that perturbations produce nonlinear ecodeficit responses, which may exhibit points of no return like asymptotes and discontinuities. The energy-based trophic chain model we derived requires minimal data and knowledge. For instance, values for assimilation and production efficiencies can be estimated from previous fieldwork, as they tend to lie in tight ranges due to fundamental constraints on energy conversion processes (Humphreys, 1979, Lindeman, 1942, Ross and Calvin, 1967). Values for population-based parameters, such as uptake rates in classic trophic chain models, do not necessarily lie in similar tight ranges. This is because they depend on the energy *Q* supplied to the ecosystem, which can exhibit considerable variation across ecosystems (e.g., about 19 times in the African savanna). Further analysis using more detailed mathematical models could illuminate the ecodeficit associated with more localized perturbations, such as specific species interactions.

To apply our framework to real-world ecosystems, we require data on population biomasses, energy inputs, and how populations use energy (e.g., for production and respiration). Our analysis of the African savanna highlights the potential of our framework to quantify the extent of ecosystem degradation and its economic implications. Our study indicates that current conservation efforts may be significantly underestimating the actual cost of restoring degraded ecosystems. Using the allometry of mass-specific metabolic rates, we estimated the monetary ecodeficit contributed just by lions is 426USD/km^2^/year, and the monetary ecodeficit contributed by all carnivores is 2, 834 USD/km^2^/year (Methods). These numbers align well with estimates between 1, 000 and 2, 000 USD/km^2^/year for the optimal operation costs for rescuing the lion population in Africa(Lindsey et al., 2018). Unfortunately, recent work reports that reserves in Africa receive between 100 and 200 USD/km^2^/year only(Lindsey et al., 2018). That is, current budgets are insufficient even to pay for the raw energy required to regenerate lion or carnivore populations, let alone the additional logistic costs for supplying that energy (e.g., workers and equipment). Indeed, the average monetary ecodeficit of 16, 401 USD/km^2^/year that we estimate to regenerate both herbivores and carnivores is about 100 times larger than current budgets, and 10 times larger than the optimal budgets previously estimated for lions only.

In line with other studies (Strassburg et al., 2020), our framework can also inform conservation priorities by identifying areas with the highest ecodeficits and guiding the allocation of limited resources. Further work improving the quality and quantity of data collection, refining embodied energy costs per location, developing more detailed dynamic models, and quantifying the effects of climate change on ecosystem energy supply would lead to better estimates of energetic ecodeficits. We believe energy can be a valuable common currency to understand and communicate better all economic costs associated with ecosystem regeneration, as it is an essential ingredient shared by economies, people, and ecosystems.

## Methods

Here we provide a summary of our methods, referring to the Supplementary Notes for details.

### Energy dynamics of a trophic chain

Let *x*_*i*_ denote the energy density of level *i* (units of energy/area). Biomass *B*_*i*_ and energy *x*_*i*_ densities are related by *x*_*i*_ = *µ*_*i*_*B*_*i*_, where *µ*_*i*_ is energy content of biomass in the trophic level *i* (units energy/mass). Typical energy contents range between 4 and 6 kcal per gram depending on the trophic level (see Table 1 in Supplementary Note 1). The energy dynamics of the trophic chain can then be derived from an energy conservation argument, assuming that the rate of change 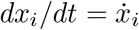 of energy density at trophic level *i* equals the assimilated energy *A*_*i*_ minus the energy that the trophic level uses for production *P*_*i*_ (e.g., predation from the next trophic level) and respiration (see Supplementary Note 1.1 for details). This results in a system of *n* = 3 differential equations parametrized by the uptake rate *α*_*i*_ ≥ 0 and energy assimilation efficiency *f*_*i*_ ≥ 0 of each level (see Eq. (S2) in Supplementary Note 1).

Let *J*_*i*_ = lim_*t→∞*_ *P*_*i*_(*t*)*/A*_*i*_(*t*) be the steady-state production efficiency of trophic level *i, i* = 1, 2. In Supplementary Note 1.2 we show that the uptake rates can be rewritten as functions of these production efficiencies as

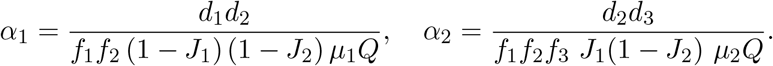

Substituting these expressions in Eq. (S2) of Supplementary Note 1 yields the following system of *n* = 3 differential equations using only energy-based parameters:

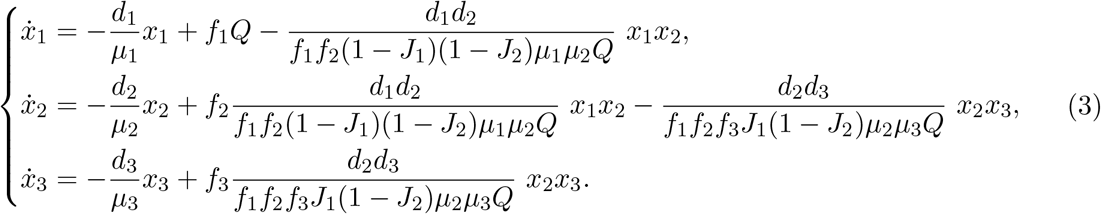

The constant external energy supply is represented by *Q*.

### Ecodeficit in model ecosystems

Solving for the interior equilibrium of Eq. (3) and using the equivalence *B*_*i*_ = *x*_*i*_*/µ*_*i*_ yields the biomass density equilibrium

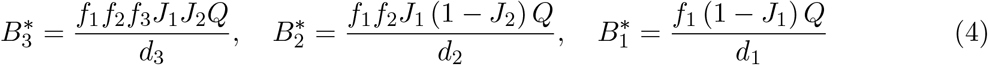

The above expressions allows us to calculate the ecodeficit factor ∆ associated to perturbations to different energetic-parameters by solving Eq. (1) of the main text as follows. Consider some arbitrary parameter *θ* to perturb as *βθ* for some constant *β*. Denote by 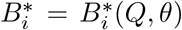 the biomass density of trophic level *i*. Then, the ecodeficit factor ∆ is the solution to the following optimization problem

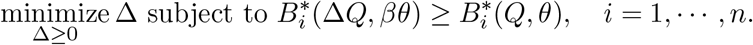

Note this optimization problem is a special case of Eq. (1) of the main text because the system has only one energy input (i.e., *m* = 1).

Supplementary Note 1.3 describes the solution of the above optimization problem to calculate the ecodeficit associated to perturbing energy-based parameters. Supplementary Note 1.4 describes the solution when perturbing population-based parameters.

### Estimating the ecodeficit in real-world ecosystems

Let *f*_*i*_, *J*_*i*_ and *d*_*i*_ denote the perturbed parameter values in a reserve, leading to the perturbed biomass densities 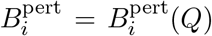 of Eq. (4). Given reference densities 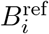, the ecodeficit factor ∆ is the minimum ∆ such that 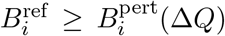. The linearity of 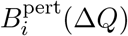 with respect to *Q* yields

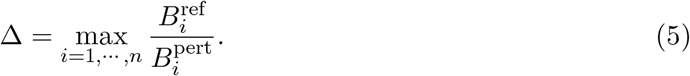

The above expression translates the problem of estimating the ecodeficit factor ∆ into estimating the reference densities 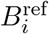.

To estimate the reference densities, consider å assumptions (i) to (iii) of the main text. Under these conditions, Eq. (4) yields

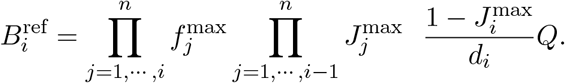

To estimate the input energy supply 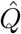, note that in our model the Net Primary Production (NPP) is *α*_1_*x*_1_*x*_2_*/µ*_2_. Evaluating this expression in the steady-state 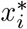 of Eq. (3) yields

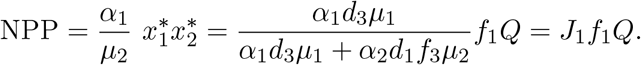

Using assumption (i) yields the estimate 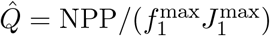 as in the main text.

Finally, we estimate the biomass-specific respiration rate 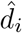 using allometry (Supplementary Note 2.1). In particular, conforming to Kleiber’s law (Kleiber, 1932), we consider the field metabolic rate *E*_*j*_ (kcal/day) of an individual of population *j* scales allometrically as 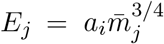. Here, 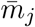 is the typical body mass of an individual of species *j*, and *a*_*i*_ is the expected field metabolic rate of an individual of trophic level *i* with unit body mass. Table 3 in Supplementary Note 2 shows estimated values for carnivora (*i* = 3) and herbivores (*i* = 2) derived in (Nagy et al., 1999). Similarly, conforming to Damuth’s rule (Damuth, 1987), we consider the expected number of individuals of population *j* per unit area *N*_*j*_ scales allometrically as 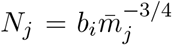, with *b*_*i*_ the expected number of individuals in one unit area. Table 4 in Supplementary Note 2 provides estimates for *b*_*i*_ for ectoterms and endoterms (Allen et al., 2002).

Substituting the estimates 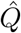 and 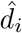 in Eq. (5) yields Eq. (2) of the main text.

### Ecodeficit contributed by specific population groups

To estimate the fraction *η* of the ecodeficit contributed by a specific group *G* of species, we consider the relative mass-specific energy consumption of the group compared to that of the whole ecosystem. Specifically, we calculate the energy consumption of organisms of population *j* via their mass-specific metabolic rate *e*_*j*_, estimated from Kleiber’s law as 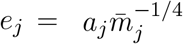 with 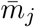 the typical body mass of individuals of population *j*. This yields 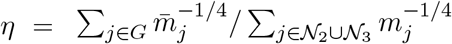 .

For the African savanna with six large carnivores and 32 large herbivores, *G* = {lion} with 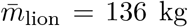 yields *η* = 0.026. For *G* = {all carnivores} we obtain *η* = 0.17. Multiplying these fractions *η* by the average monetary ecodeficit of 16, 401 USD/km^2^/year yields estimates of 426 USD/km^2^/year for lions, and 2,834 USD/km^2^/year for all carnivores.

## Acknowledgments

S.S. acknowledges support from the National Science Foundation under Grant No. DEB-2436069 and MIT Google Program for Computing Innovation. M.T.A. acknowledges the financial support provided by CONACyT grant No. A1-S-13909.

## SUPPLEMENTARY NOTES

### 1. A trophic chain model based on energy flow and conservation

#### 1.1. Derivation of the model

We consider four trophic levels as follows: level 0 corresponds to basal resources, level 1 to primary producers such as plants, level 2 to primary consumers such as herbivores, en level 3 to secondary consumers such as predators. For each trophic level *i*, we denote

- *B*_*i*_ its biomass density in kg/km^2^.
- *x*_*i*_ its energy density in kcal/km^2^.

Let *µ*_*i*_ *>* 0 denote the energy content per biomass kilogram of level *i* (units kcal/kg). Note that *x*_*i*_ = *µ*_*i*_*B*_*i*_. Time is denoted by *t* (units years). Units are written in gray color to the right of equations. The model is based in the following assumptions.

##### For trophic level 0

- Let

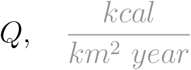

denote the supply rate of energy entering into the ecosystem.
- As in classic models of trophic chains [1], let *α*_0_*B*_0_ be the **resource uptake** (amount of resource *B*_0_ consumed by unit of biomass density *B*_1_ per unit time). Units are *α*_0_ [1/year] [1/unit of *B*_1_] and *B*_0_ [kg/km2]. The corresponding **energy uptake** is obtained multiplying by the energy content *µ*_0_ of the basal resource,

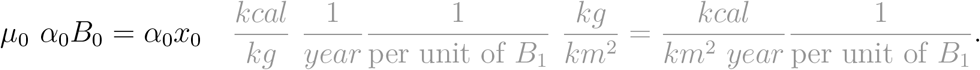
- The above uptake is per unit of biomass density *B*_1_. Therefore, the **total energy uptake** is obtained multiplying by *B*_1_

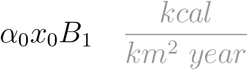 Note the above expression has the usual form for power [2], that is, power = thermodynamic flux *×* thermodynamic force.
- Energy conservation implies that the change 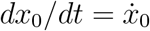 in energy at level 0 equals the input energy minus the energy uptake,

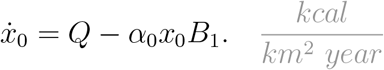

Using the equivalence *x*_0_ = *µ*_0_*B*_0_ above equation can be rewritten in terms of biomass density only as

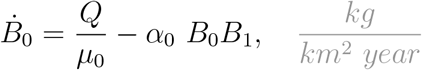

or in terms of energy density only using *B*_1_ = *x*_1_*/µ*_1_ as

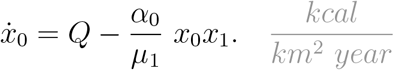

##### For trophic level 1

- From the total energy uptake *α*_0_*x*_0_*B*_1_, only a fraction *κ*_1_ ∈ [0, 1) is assimilated by the trophic level. Energy conservation implies that the **change** 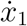 **in energy** at level 1 equals this assimilated energy minus losses due to respiration *R*_1_ and predation *P*_1_,

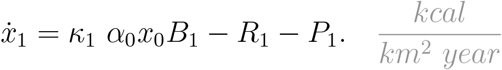
- We assume *R*_1_ = *d*_1_*B*_1_, where

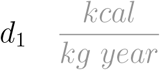

is the mass-specific respiration of the trophic level.
- Let *α*_1_*B*_1_ be the resource uptake rate that trophic level 2 makes on trophic level 1 (per unit of biomass *B*_2_). Units are *α*_1_ [1/year] [1/unit of *B*_2_] and *B*_1_ [kg/km2]. The corresponding **energy uptake** is obtained multiplying by *µ*_1_,

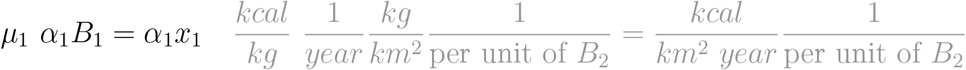 The total energy uptake *P*_1_ is obtained multiplying by *B*_2_

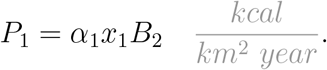
- From the above, energy conservation implies

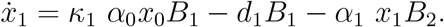

Using the equivalences *x*_0_ = *µ*_0_*B*_0_, *x*_1_ = *µ*_1_*B*_1_, and *x*_3_ = *µ*_3_*B*_3_, the above equation can be rewritten in terms of biomass density only as

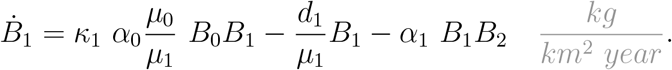

or in terms of energy density only

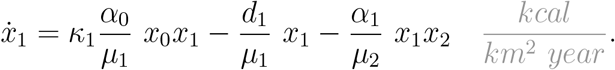

##### For trophic level 2

- Only a fraction *κ*_2_ ∈ [0, 1) of the total energy uptake *α*_1_*x*_1_*B*_2_ is assimilated. Energy conservation implies that the change 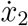 in energy at level 2 equals this assimilated energy minus loses due to respiration *R*_2_ and predation *P*_2_,

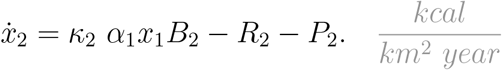
- We assume *R*_2_ = *d*_2_*B*_2_, where

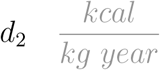

is the mass-specific respiration of the trophic level.
- Let *α*_2_*B*_2_ be the resource uptake rate that trophic level 3 makes on trophic level 2 (per unit of biomass *B*_3_). Units are *α*_1_ [1/year] [1/unit of *B*_3_] and *B*_2_ [kg/km2]. The corresponding **energy uptake** is obtained multiplying by *µ*_2_,

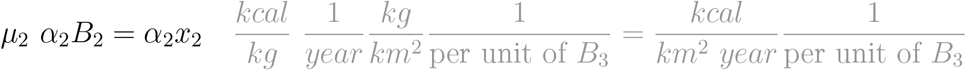 The total energy uptake *P*_2_ is obtained multiplying by *B*_3_

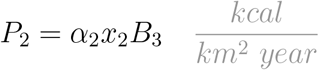
- From the above, energy conservation implies

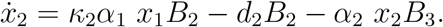

Using the equivalence *x*_*i*_ = *µ*_*i*_*B*_*i*_ we can rewrite the above equations in terms of biomass density only

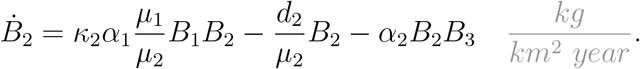

or in terms of energy density only

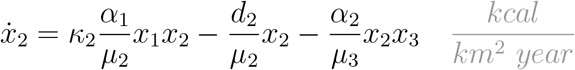

##### For trophic level 3

- Only a fraction *κ*_3_ ∈ [0, 1) of the total energy uptake *α*_2_*x*_2_*B*_3_ is assimilated. Energy conservation implies that the change 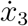 in energy at level 3 equals this assimilated energy minus loses due to respiration *R*_3_

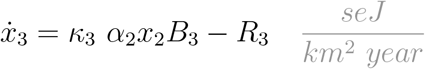
- We assume *R*_3_ = *d*_3_*B*_3_, where

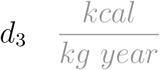

is the mass-specific respiration of the trophic level.
- From the above we conclude

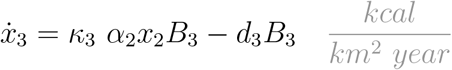

Using the equivalence *x*_*i*_ = *τ*_*i*_*µ*_*i*_*B*_*i*_, we can rewrite this equation in terms of biomass density only as

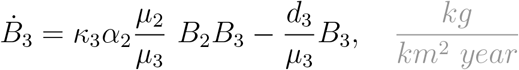

or using *B*_*i*_ = *x*_*i*_*/τ*_*i*_*µ*_*i*_ in terms of energy density only as

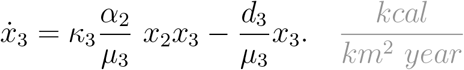

To summarize, the population dynamics in terms of biomass density *B*_*i*_ (units kg/km2) reads

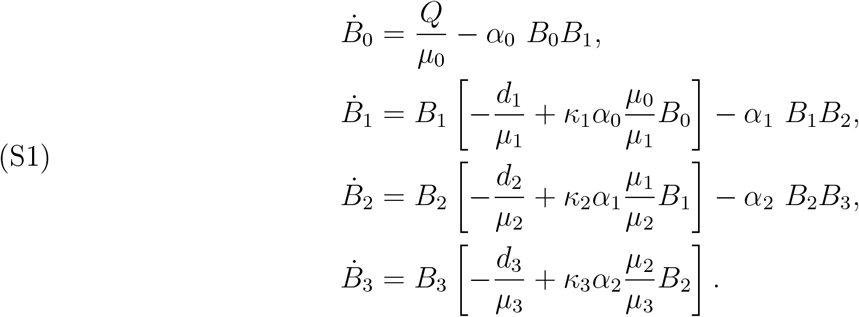

**Table 1.** Typical energy parameters. Derived from Odum pp. 109. Efficiency is the product of utilization and asimilation efficiencies. Production is the fraction of the energy used for production, the rest used for respiration. Energy content is derived from Table 3-1.

| trohic level | efficiency ($f_i$) | production ($J_i$) | energy content ($\mu_i$) |
| --- | --- | --- | --- |
| units | adimensional | adimensional | kcal/g Carbon |
| 1. plants | [0.006, 0.007] | [0.6, 0.7] | 4.5 |
| 2. herbivores | [0.14, 0.2] | [0.05, 0.1] | 5.6 |
| 3. predator | [0.27, 0.38] | [0.05, 0.1] | 5.6 |

Equivalently, the population dynamics in terms of energy density *x*_*i*_ (units kcal/km2) reads

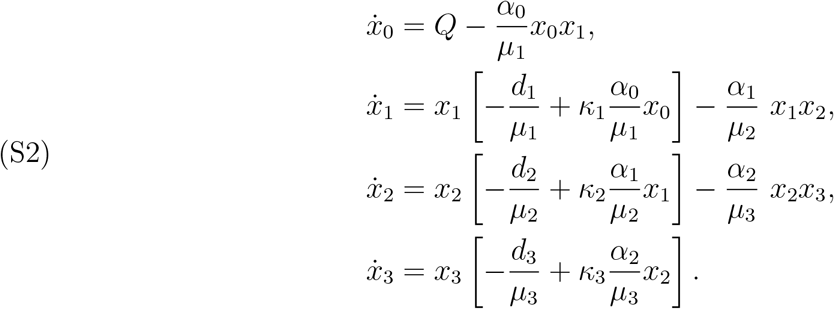

Note that it is rather difficult to set plausible ranges for the values of the parameters *α*_*i*_ of the model. To circumvent this limitation, we next explain how to re-parametrize the model in terms of energy efficiency parameters.

#### 1.2. Parametrization based on energy efficiency and use

To parametrize the model we consider the steady-state energy efficiency *f*_*i*_ and fraction of assimilated energy used for production *J*_*i*_ for trophic level *i*. Typical values for these parameters are reported by Odum as shown Table 1.

Because the parameters *κ*_*i*_ represent energy assimilation efficiency, we can immediately conclude that *κ*_*i*_ = *f*_*i*_. Therefore, the objective is to rewrite the parameters *α*_*i*_ in terms of the parameters *J*_*i*_.

With this objective, from (S2) in equilibrium we deduce

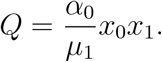

Next, note that the Gross Primary Production (GPP) in the model is

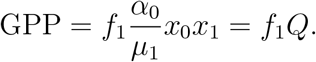

Consider now the Net Primary Production (NPP). Since GPP = NPP + Respiration, we obtain

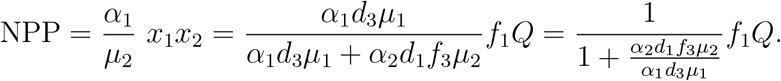

Because GPP = *f*_1_*Q*, the above expression implies that the fraction *J*_1_ of energy used for production satisfies

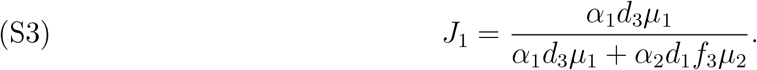

At the next trophic level, the assimilated energy is *f*_2_NPP. Consider now the Secondary Production (SP). From the condition *f*_2_NPP = SP + Respiration, we deduce the secondary production is

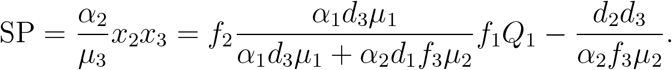

The fraction *J*_2_ of energy used for production is *J*_2_ = SP*/*(*f*_2_NPP). We thus deduce

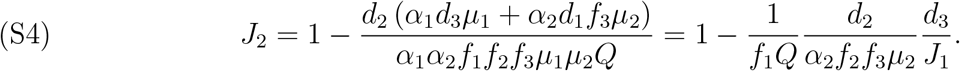

Using the above equations, we can express *α*_1_ and *α*_2_ as functions of *J*_1_ and *J*_2_ as follows:

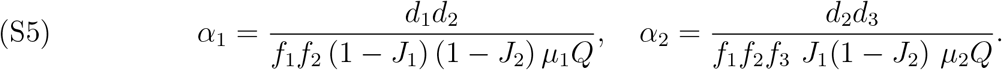

Using the values (*f*_*i*_, *J*_*i*_, *µ*_*i*_) from Table 1, the only parameters missing for the model are the respiration rates *d*_*i*_ of trophic levels, and the energy supply input *Q*.

#### 1.3. The ecodeficit of perturbing energetic-based parameters

Solving for the positive equilibrium energy densities 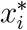 in Eq. (S2) yields

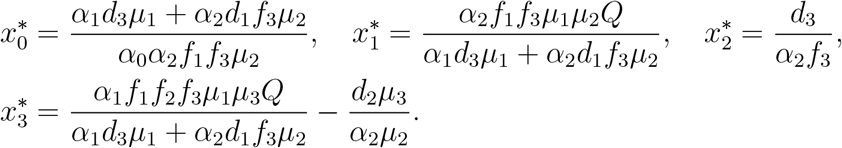

Using the equivalence *B*_*i*_ = *x*_*i*_*/µ*_*i*_ for biomass density *B*_*i*_ and rewriting in terms of assimilation *f*_*i*_ and production *J*_*i*_ efficiencies yields

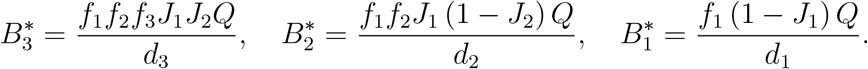

The above expressions allows us to rewrite the biomass densities of the first two levels as functions of the biomass density of the last trophic level:

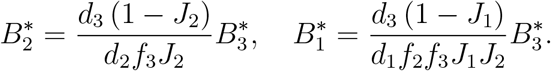

The above expressions allows us to calculate the ecodeficit factor ∆ associated to perturbations to different energetic-parameters as follows. Consider some arbitrary parameter *θ* to perturb as *βθ* for some constant *β*. Denote by 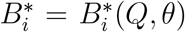 the biomass density of trophic level *i*. Then, the ecodeficit factor ∆ is the solution to the following optimization problem

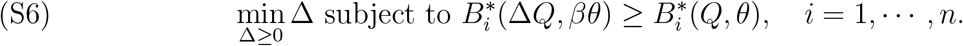

To illustrate the process of calculating the ecodeficit, consider a perturbation to mass-specific respiration rate *d*_1_ ↦ *βd*_1_. The ecodeficit factor is the minimum ∆ *≥*0 such that the following inequalities are satisfied:

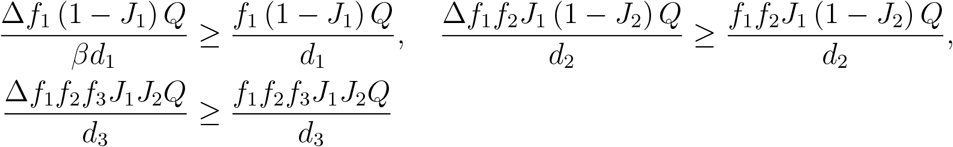

The above inequalities can be simplified to

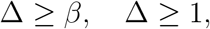

from which we deduce the ecodeficit factor is ∆ = max*{β*, 1*}*.

Similar calculations shows the following:

1. The ecodeficit associated to perturbing a respiration rate *d*_*i*_ ↦ *βd*_*i*_ is

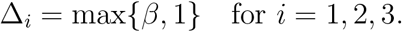
2. The ecodeficit associated to perturbing the first assimilation efficiency *f*_1_ ↦ *βf*_1_ is

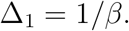 The ecodeficit associated to perturbing the assimilation efficiency *f*_*i*_ *↦βf*_*i*_ of the second and third levels is

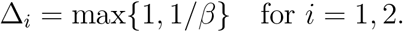
3. The ecodeficit associated to perturbing the production efficency *J*_*i*_ → *βJ*_*i*_ is

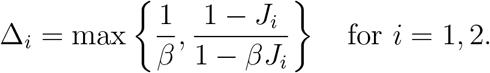

Note that for small perturbations *β ≈* 1, the ecodeficit associated to perturbing respiration rates is *β*, and the ecodeficit of perturbing assimilation or production efficiencies is proportional to 1*/β*. That is, in this case the ecodeficit can be calculating as the condition under which the last trophic level reaches the reference value.

#### 1.4. The ecodeficit of perturbing population-based parameters

In terms of the mortality rates *m*_*i*_ = *d*_*i*_*/µ*_*i*_ and uptake rates *α*_*i*_, the positive equilibrium of biomass densities 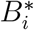 are

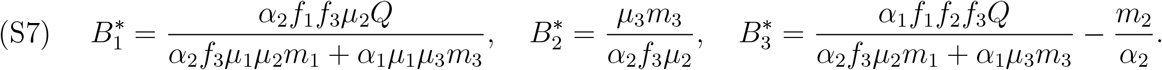

Solving the optimization problem defining the ecodeficit for each parameter yields:

1. The ecodeficit associated to perturbing the uptake rate *α*_1_ ↦*βα*_1_ of the first trophic level is

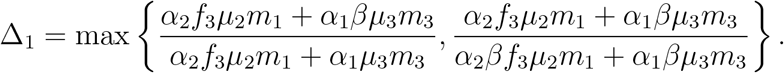 The ecodeficit ∆_2_ of the second trophic level exist only if the perturbation reduces its uptake rate (i.e., *β≤* 1). This happens because the biomass density of the second trophic level 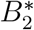 does not directly depend on *Q*. Under this condition, the ecodeficit is

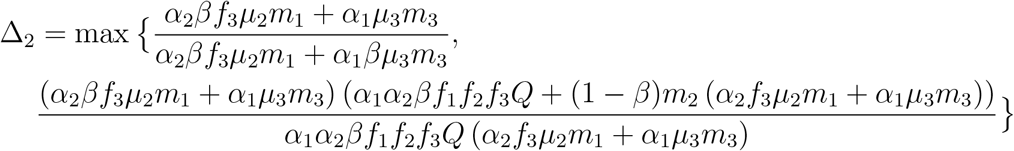
2. The ecodeficit associated to perturbing the first mortality rate is

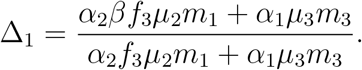 When perturbing the second mortality rate

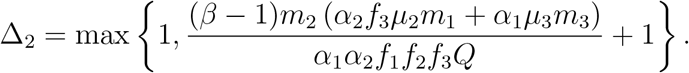 When perturbing the third mortality rate, the ecodeficit is finite only if *β ≥*1 (i.e., if mortality increases). In such case, the ecodeficit is

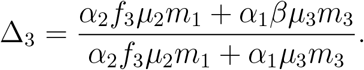

### 2. Ecodeficit for large mammales in the African savanna

#### 2.1. Estimating mass-specific respiration rates

To calculate the reference biomass densities, we estimate the expected value for the biomass-specific respiration rate *d*_*i*_ (units kcal/kg/-day) based on allometry as follows:

1. We quantify the energy spent by an organisms using its field metabolic rate (units kcal/day). The field metabolic rate represents the total energy expenditure of a free-living animal in its natural environment, encompassing all its daily activities, including foraging, moving, and maintaining bodily functions, essentially measuring the total energy an animal uses in the wild throughout a day. For herbivores (trophic level *i* = 2) and carnivores (trophic level *i* = 3), we can estimate the field metabolic rate *E*_*j*_ that one individual of population *j* requires as a function of its typical body mass 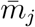 [3]:

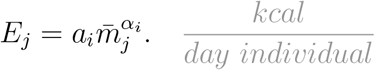 Above, the parameters *a*_*i*_ and *α*_*i*_ are the allometric coefficient and exponent for trophic level *i*, respectively. Values for these parameters are provided in Table 3.
2. The expected number *N*_*j*_ of individuals of population *j* per unit area can also be estimated from allometry using Damuth’s rule [4, 5]:

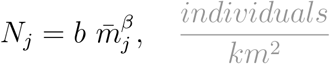 The parameters *b* and *β* are the allometric coefficient and exponent, with values provided in Table 4. Because there is no significant difference between these value for carnivores and herbivores, we use the same value for both trophic levels.

To calculate the respiration rate per unit biomass of trophic level *i*, consider a reserve of area *A*. Let 𝒩_*i*_ denote the set of all populations belonging to trophic level *i*. For a population *j* ∈*𝒩*_*i*_, we expect a total number of *AN*_*j*_ individuals. Each of those individuals will spend *E*_*j*_ kcal per day. Therefore, the total energy spent by all individuals in the trophic level is

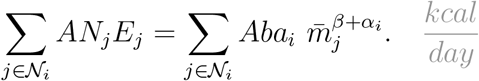

On the other hand, the total biomass expected from allometry is the number *AN*_*j*_ of individuals of population *j*, multiplied by their typical body mass 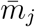:

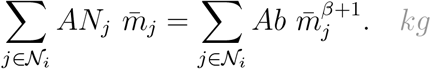

Finally, the biomass-specific respiration rate (kcal/kg/day) expected by allometry can be derived from the quotient of the last two quantities:

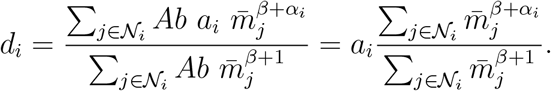

**Table 2.**
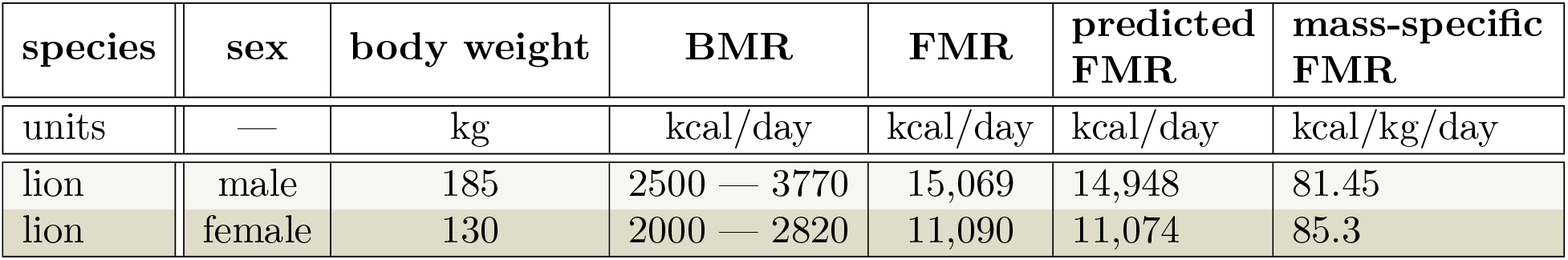
Basal Metabolic Rate (BMR) and Field Metabolic Rate (FMR) for lions. Data from [6]. The FMR is predicted using alometric scaling from Naggy (see Table 3).

| species | sex | body weight | BMR | FMR | predicted FMR | mass-specific FMR |
| --- | --- | --- | --- | --- | --- | --- |
| units | — | kg | kcal/day | kcal/day | kcal/day | kcal/kg/day |
| lion | male | 185 | 2500 — 3770 | 15,069 | 14,948 | 81.45 |
| lion | female | 130 | 2000 — 2820 | 11,090 | 11,074 | 85.3 |

**Table 3.**
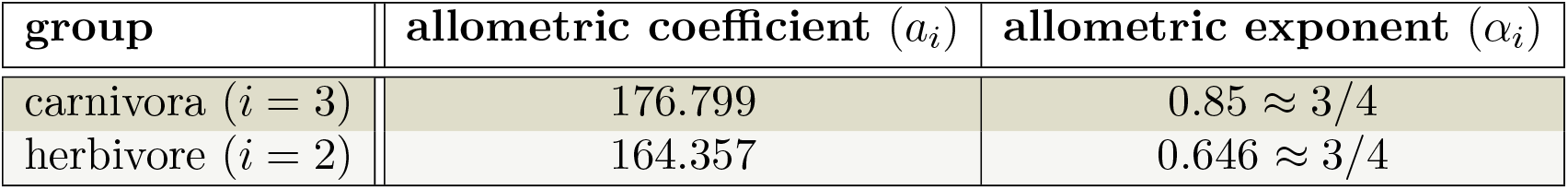
Allometric estimation of Field Metabolic Rate derived from Nagy [3]. The equation is 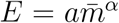 where *E* is the field metabolic rate (kcal/day) and *m* is the typical body mass of an individual (kg).

| group | allometric coefficient ($a_i$) | allometric exponent ($\alpha_i$) |
| --- | --- | --- |
| carnivora ($i = 3$) | 176.799 | $0.85 \approx 3/4$ |
| herbivore ($i = 2$) | 164.357 | $0.646 \approx 3/4$ |

**Table 4.** Allometric estimation for the density of individuals [5]. The equation is 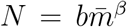 where *N* is the the number of individuals per km^2^ and *m* is the typical body mass of an individual (kg).

| group | allometric coefficient ($b$) | allometric exponent ($\beta$) |
| --- | --- | --- |
| ectotherms and endotherms | $e^{-19.63} = 2.984 \times 10^{-9}$ | $-0.78 \approx -3/4$ |

